# Chromoprotein-modified plant pathogenic bacteria: tools for experimental tracking and visualization

**DOI:** 10.1101/2025.08.26.672430

**Authors:** Lindsey Burbank

**Affiliations:** USDA, Agricultural Research Service, San Joaquin Valley Agricultural Sciences Center, Parlier, CA

**Keywords:** chromoproteins, *Xylella fastidiosa*, *Pseudomonas syringae*, *Xanthomonas campestris*, *Pantoea stewartii*, plant pathogens

## Abstract

Microbiology research often requires tracking of specific bacterial strains within a host infection system or in the environment, as well as differentiation of strains in a co-infection or microbe-microbe interaction scenario. Various tools are used for this purpose including antibiotic resistance marker genes, fluorescent proteins, DNA sequence-based methods, and phenotypic markers. Chromoproteins produce intense pigmentation visible in ambient light, and are a unique option for bacterial tracking that does not require use of antibiotics, specialized equipment, or DNA sequencing. Development of traceable bacterial strains across a wide range of species is important to facilitate the investigation of challenging research questions and expand our understanding of microbial dynamics in complex environments. In this study, different species of plant pathogenic bacteria (*Xylella fastidiosa, Pantoea stewartii, Pseudomonas syringae* and *Xanthomonas campestris*) were modified with a set of chromoproteins and tested in plant infection assays to evaluate chromoprotein stability and impact on bacterial pathogenicity. Chromoprotein modification by chromosomal insertion was highly successful in *X. fastidiosa*, and stable during infection in grapevines. Plasmid-based expression of chromoproteins in *P. stewartii, P. syringae*, and *X. campestris* had mixed results depending on the specific species-chromoprotein combination. Overall, these results provide some successful chromoprotein-modified plant pathogen strains for use by the research community, as well as insight into which chromoproteins might be best utilized in different bacterial species.

**Importance:** Investigation of challenging research questions in plant health requires availability of the necessary microbial tools. This study adapts chromoprotein-modification for bacterial tracking of plant pathogens during host infection and plant-plant transmission. Successful development of visibly colored strains of *Xylella fastidiosa*, *Pantoea stewartii*, *Pseudomonas syringae*, and *Xanthomonas campestris* provides a useful resource for research and educational purposes, as well as insight into optimization of chromoprotein expression in a range of plant pathogen species.

## Introduction

Traceability of specific bacterial strains and mutants is an essential component of experimental design when testing microbe-microbe, microbe-host, and microbe-environment interactions. Antibiotic resistance genes, fluorescent proteins (1, 2), and other distinctive genetic (3) or phenotypic markers (4) can be used for this, but each strategy has unique tradeoffs (5–7). Antibiotic markers have limitations in some bacterial species and environmental scenarios, such as existence of background levels of similar genes in wild microbes, or concerns about contamination of water or soil (8, 9). Fluorescent proteins require specialized equipment for imaging and visualization (10), and can be subject to interference from chemical exposure or host autofluorescence (11–13). Sequence or DNA barcode-based approaches can require a significant amount of sequencing which is costly at scale and not practical across the whole spectrum of resource availability (14, 15). Chromoproteins, highly pigmented proteins often derived from corals and marine animals (16, 17), can be a useful alternative for research experiments, or education and outreach demonstrations (18–23). These proteins produce visible colors in ambient light which enables tracking specific bacterial strains through an environmental system, or visualizing interactions of different strains without imaging equipment. Chromoproteins are available in a wide range of color profiles, and can be optimized for individual research systems or for expression in specific bacterial species (24).

The goal of this study is to test chromoprotein expression in plant-pathogenic bacteria as a tool for rapid tracking and differentiation of bacterial strains in plant disease systems. The primary focus is on creating *Xylella fastidiosa* strains with stable chromoprotein expression during plant infection, with some additional exploration of plasmid-based chromoprotein expression in *Pantoea stewartii*, *Pseudomonas syringae*, and *Xanthomonas campestris*. *Xylella fastidiosa* is a pathogen that presents unique research and disease mitigation challenges due to its fastidious nature, non-specific insect transmission, and broad plant host-range in woody perennial plants (25, 26).

Availability of multiple chromoprotein-tagged strains of *X. fastidiosa* would facilitate investigation of plant-plant transmission patterns, interaction of different strains and subspecies in the plant host, and within-plant movement under different conditions, all of which are important research questions to improve disease mitigation strategies (27). The bacterial genera *Pantoea*, *Pseudomonas*, and *Xanthomonas*, contain a wide range of important plant pathogen species, plant disease model systems, and beneficial microbes that are used extensively in agriculture and plant science research (28–37). This study used *P. stewartii*, *P. syringae*, and *X. campestris* species to test expression of chromoproteins carried on a broad host range plasmid during in vitro growth and plant infection. Overall, success varies based on the bacterial species, specific chromoprotein, and plant infection dynamics, but these results lay groundwork for continued improvement of chromoprotein-modified plant pathogen strains for a variety of research purposes. Additionally, as plant pathogens are generally safe and easy to work with in teaching and outreach activities, colored bacterial strains could be used in novel ways for demonstration and educational purposes.

## Materials and methods

### Bacterial strains and general culture conditions

All cloning and plasmid manipulation steps were done in *Escherichia coli* Top10 (ThermoFisher Scientific), and any mention of *E. coli* in the methodology refers to this strain. *Xylella fastidiosa* strains used in this study were *X. fastidiosa* subsp. *fastidiosa* M23 (38) and *X. fastidiosa* subspecies *fastidiosa* Temecula-1 (39). *Pantoea stewartii* strain used was *P. stewartii* subsp. *stewartii* DC283 (40), *Pseudomonas syringae* strain used was *P. syringae* pv tomato DC3000 (41), and *Xanthomonas campestris* strain was *X. campestris* pv *campestris* 8004 (42). *E. coli* was cultured in LB medium with addition of the following antibiotics when necessary: spectinomycin (100 µg/mL), gentamicin (10 µg/mL), kanamycin (30 µg/mL). *Xylella fastidiosa* was cultured in PD3 medium (43) supplemented with gentamicin (5 µg/mL) when necessary. *P. stewartii*, *X. campestris*, and *P. syringae* were cultured on tryptic soy agar (TSA) with gentamicin (10 µg/mL), nalidixic acid (30 µg/mL) or rifampicin (50 µg/mL) when necessary.

### Chromoprotein modification of *Xylella fastidiosa*

The pIDMv5K plasmid series (Bionomica Labs) was used as a source of six different chromoproteins (Table 1, Fig. 1A). Chromoprotein genes with promoter and terminator elements included were PCR-amplified from pIDMv5K source plasmids using primers pIDM-fwd/pIDM-rev (Table 2). Amplicons were cloned into pCR8/GW/TOPO vector following manufacturer’s instructions and plated on selective plates with spectinomycin. Colonies with visible color expression were chosen and plasmids were extracted using a Qiaprep miniprep kit (Qiagen). Fifty ng of pCR8-chromoprotein clones were recombined with 150 ng of pAX1-Gm-GW (44, 45) vector using LR Clonase II (ThermoFisher) following the manufacturers instructions. Clones of pAX1-chromoprotein with visible color expression were chosen for sequencing. Whole plasmid sequencing was performed by Plasmidsaurus Inc. Full plasmid sequences were submitted to GenBank under accession numbers included in Table 1. For chromosomal insertion in *X. fastidiosa*, pAX1-chromoprotein constructs were transformed into *X. fastidiosa* strains M23 and Temecula-1 by natural transformation as follows. *X. fastidiosa* strains were grown from frozen glycerol stocks on PD3 plates for one week at 28°C. Cells were scraped off plates and resuspended in liquid PD3 medium at a cell density of OD_600nm_ = 0.25. Ten uL of cell suspension was combined with 300 ng of plasmid DNA, spotted onto PD3 plates, and allowed to dry. Then plates were incubated at 28°C for 4 days. After initial incubation, each spot of cells was scraped off plates and resuspended in 500 µL of PD3 medium, then plated on PD3 with gentamicin for selection of transformants. Each transformation included a negative control (no DNA added) and a positive control (plasmid pAX1-Gm with no insert). Transformant colonies appeared after 10-14 days incubation at 28°C, and transformants with visible color were selected for confirmation and further experiments. Modified strains were confirmed as *X. fastidiosa* by PCR with primers RST31/RST33 (Table 2), and for insertion of the chromoprotein gene in the correct location with primers NS1-fwd/NS1-rev (Table 2).

**Figure 1.**
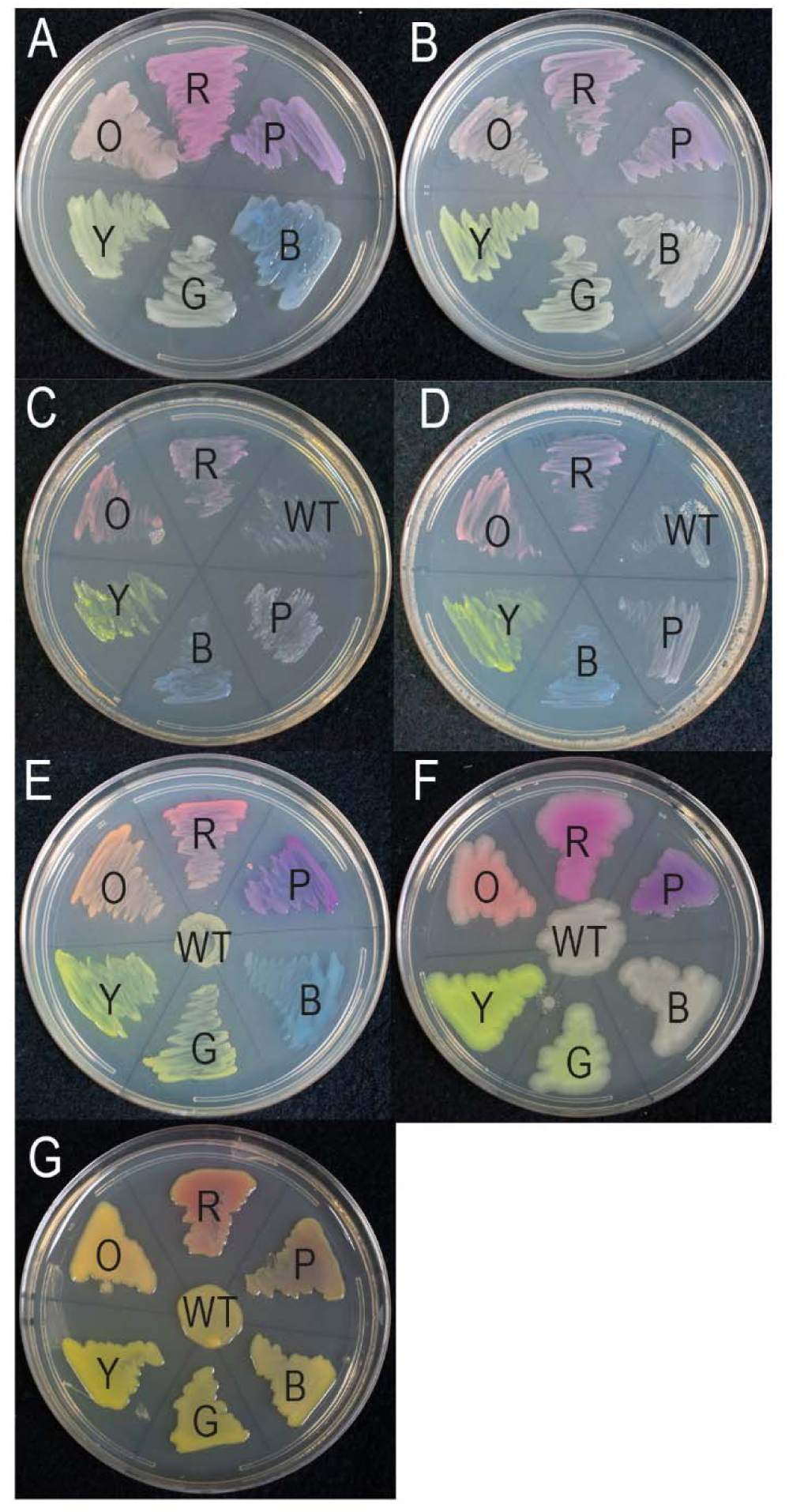
Chromoprotein-modified strains of plant pathogenic bacteria. Chromoprotein genes were cloned from the pIDMv5K plasmid series for expression in *Xylella fastidiosa*, *Pantoea stewartii*, *Pseudomonas syringae*, and *Xanthomonas campestris*. **A)** *Escherichia coli* with pIDMv5K series plasmids containing chromoproteins after 24 hours growth at 37°C. **B)** *E. coli* with low copy, broad host range plasmid pBBR5pemIK expressing chromoproteins after 24 hours at 37°C. **C)** *Xylella fastidiosa* strain Temecula-1 and **D)** strain M23 with chromosomal insertions of chromoproteins after 7 days growth at 28°C. **E)** *Pantoea stewartii*, **F)** *Pseudomonas syringae*, and **G)** *Xanthomonas campestris* with chromoproteins expressed on plasmid pBBR5pemIK after 3 days of growth at 28°C. WT = wild type, R = red, O = orange, Y = yellow, G = green, B = blue, P = purple.

**Table 1.** Plasmids used in this study.

| Plasmid | Description | Sequence | Source reference |
| --- | --- | --- | --- |
| pIDMv5K-RED | Source of red chromoprotein eforCP, Kan<sup>R</sup> | Bionomica Labs (58) | Bionomica Labs (58) |
| pIDMv5K-ORANGE | Source of orange chromoprotein YukonOFP, Kan<sup>R</sup> | Bionomica Labs (58) | Bionomica Labs (58) |
| pIDMv5K-YELLOW | Source of yellow chromoprotein DasherGFP, Kan<sup>R</sup> | Bionomica Labs (58) | Bionomica Labs (58) |
| pIDMv5K-GREEN | Source of green chromoprotein fuGFP, Kan<sup>R</sup> | Bionomica Labs (58) | Bionomica Labs (58) |
| pIDMv5K-BLUE | Source of blue chromoprotein aeBlue, Kan<sup>R</sup> | Bionomica Labs (58) | Bionomica Labs (58) |
| pIDMv5K-PURPLE | Source of purple chromoprotein tsPurple, Kan<sup>R</sup> | Bionomica Labs (58) | Bionomica Labs (58) |
| pCR8/GW/TOPO | PCR cloning vector compatible with Gateway cloning, Sp<sup>R</sup> |  | Thermo Fisher Scientific |
| pBBR5pemIK | Broad host range vector with toxin-antitoxin system added for stability, Gm<sup>R</sup> | GenBank: KX036764.1 | (46) |
| pBBR5pemIK-GW |  | GenBank: KX036764.1 | (46) |
| pAX1-Gm-GW |  | PV932842 | (45) |
| pBBR5pemIK-RED |  | PV983734 | This study |
| pBBR5pemIK-ORANGE |  | PV983735 | This study |
| pBBR5pemIK-YELLOW |  | PV983736 | This study |
| pBBR5pemIK-GREEN |  | PV983737 | This study |
| pBBR5pemIK-BLUE |  | PV983738 | This study |
| pBBR5pemIK-PURPLE |  | PV983739 | This study |
| pAX1-RED |  | PV932843 | This study |
| pAX1-ORANGE |  | PV932844 | This study |
| pAX1-YELLOW |  | PV932845 | This study |
| pAX1-BLUE |  | PV932846 | This study |
| pAX1-PURPLE |  | PV932847 | This study |

**Table 2.**
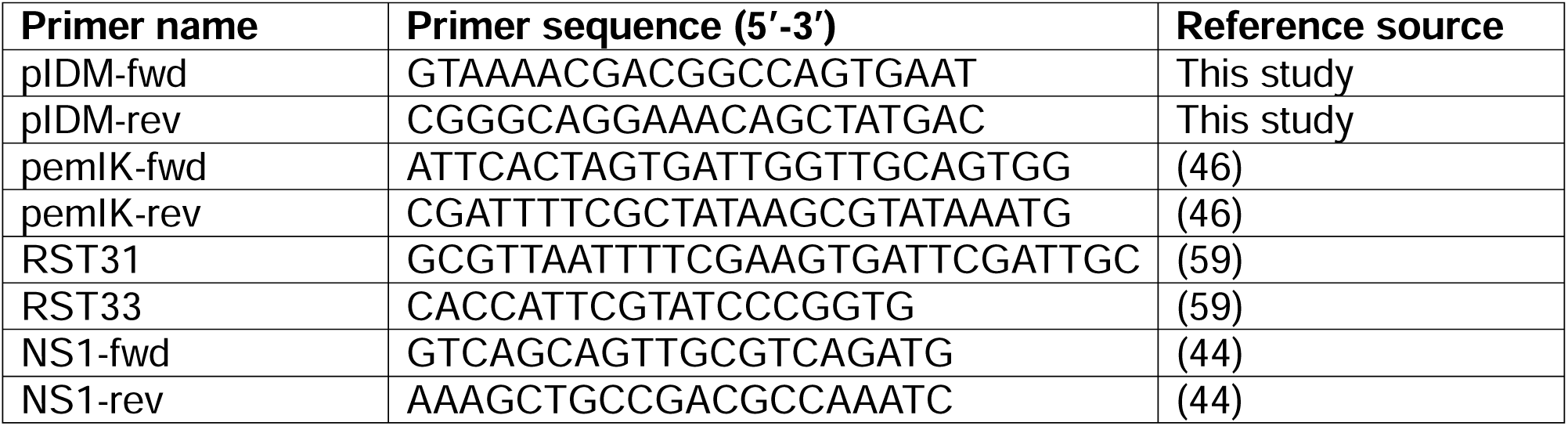
Primer sequences.

### Chromoprotein modification of Pantoea stewartii, Pseudomonas syringae, and Xanthomonas campestris

Plasmid-based chromoprotein expression was used for *P. stewartii*, *P. syringae*, and *X. campestris* with the stable, broad-host range plasmid vector pBBR5pemIk (46). Chromoprotein plasmid constructs were created by recombination of pCR8-chromoprotein plasmids described above with the pBBR5pemIK-GW vector using LR Clonase II (ThermoFisher) following the manufacturer’s instructions. Clones with visible color expression were chosen for sequencing. Whole-plasmid sequencing was performed by Plasmidsaurus Inc. Full plasmid sequences were submitted to GenBank under accession numbers included in Table 1. Color expression pBBR5pemIK plasmids (Fig. 1B) were transformed into *P. stewartii*, *P. syringae*, and *X. campestris* using electroporation as follows. Cells were grown in TSA medium to a cell density of OD_600nm_ = 0.3 and harvested by centrifugation of 2 mL aliquots at 9000 rpm for 3 minutes. Cell aliquots were washed twice in sterile distilled H_2_O and twice in 10% glycerol by vortex-mixing to resuspend, centrifugation at 9000 rpm for 3 minutes, and removal of supernatant. After washing, cell pellets were resuspended in 100 µL of 10% glycerol and either stored at -80°C or immediately used for transformation. For electroporation, 150-200 ng of plasmid DNA was mixed with 100 µL of cells and kept on ice for 5 minutes. Plasmid-cell mixture was transferred to a 0.2 cm sterile electroporation cuvette (BioRad) pre-chilled on ice and electroporated using a GenePulser XCell instrument (BioRad) with the electrical settings as follows: 2.5 kV (12.5 kV/cm), 25 µF, 200 Ω. Immediately following electroporation, 1 mL of TSA medium was added to the cells which were then transferred to a microfuge tube and incubated on a shaker for 1 hour at 28°C and 180 rpm shaking. Cells were then plated on TSA medium supplemented with gentamicin for selection and incubated at 28°C for 48 hours. Transformation negative controls were cells electroporated with no DNA and empty vector pBBR5pemIK was also transformed with each bacterial species. Colonies with color expression were selected for further screening. In some cases, if a significant number of transformants were present but colors were not easily visible due to the presence of naturally occurring pigments or low color expression in some species, transformants were further screened with PCR to confirm presence of the plasmid (primers pemIK-fwd/pemIK-rev).

### Plant inoculations with *Xylella fastidiosa*

Grapevines (*Vitis vinifera* cv Cabernet Sauvignon) were grown in 3.8L pots in Sunshine Mix #1 professional growing mix (SunGro Horticulture) in a temperature-controlled greenhouse maintained at 24-37°C with natural light. Plants were grown from rooted cuttings until the vine reached 1 meter length before inoculation. Plants were inoculated with *X. fastidiosa* following the pin-prick inoculation protocol as previously described (47). Briefly, *X. fastidiosa* was grown on PD3 plates for 5 days at 28°C. Cells were scraped off plates in 1XPBS and diluted to a concentration of OD_600nm_=0.25 (∼10^8^cfu/mL). Two 20 µL drops of cell suspension were placed on the side of the grapevine stem and pierced with a 20-gauge needle to allow uptake directly into the xylem. Inoculation was repeated on the opposite side of the stem for each plant to receive a total of 80 µL of cell suspension. For plants inoculated with two different strains, half the inoculum came from each strain. Negative control plants were mock inoculated with 1XPBS. Three plants were inoculated with each strain or strain combination. After inoculation, plants were then maintained in the greenhouse until disease symptoms were visible. Petiole samples for bacterial isolation were collected from each plant once it reached a disease score of 2-3 on a previously described 0-5 disease rating scale (47). All inoculated plants reached this level of disease between 8-12 weeks post-inoculation. Three petioles were collected from symptomatic leaves on each plant and processed individually for bacterial isolation.

Petioles were weighed, then surface sterilized by placing in 70% ethanol for 1 minute, 20% bleach for 1 minute, and then washed in sterile water twice for 1 minute each wash. Sterilized petioles were ground in plastic mesh grinding bags (Agdia Inc) with 2 mL 1XPBS. Serial dilutions of buffer containing ground plant material were made in 1XPBS and then plated on PD3 plates. Plates were incubated at 28°C for 14 days for colonies to be large enough for counting, and for visible color expression.

### Plant inoculations with *Pantoea stewartii*

Sweet corn (*Zea maize* cv Jubilee) was grown from seed in 10 cm pots of Sunshine Mix #1 professional growing mix (SunGro Horticulture) in a temperature-controlled greenhouse maintained at 24-37°C with natural light. Seedlings were inoculated when they were 14 days old, and maintained in the greenhouse throughout the experiments. For inoculation, bacterial strains were grown on TSA plates for 48 hours at 28°C. Liquid cultures of 4 mL TSA medium were inoculated with single colonies and grown overnight for cells to reach a concentration of OD_600nm_=1.0. Cells were centrifuged at 9000 rpm for 3 minutes, and cell pellets were washed twice in 1XPBS. After washing, cell pellets were resuspended to the original volume in 1XPBS with 0.2% Tween 20. 150 µL of cell suspension was inoculated into the whorl of the leaves, with four plants inoculated per bacterial strain and four plants mock inoculated with buffer. Four days after inoculation, the largest leaf was collected from 3-4 plants per treatment for bacterial isolation. A 10 cm section of the leaf was cut out, weighed, and surface sterilized in 70% ethanol for 30 seconds, 20% bleach or 30 seconds, and then washed twice for 1 minute in sterile water. Leaf sections were then transferred to a sterile grinding bag (Agdia) and crushed in 2 mL of 1XPBS. Buffer containing crushed plant material was serially diluted in 1XPBS and plated on TSA medium. After 48 hours of incubation at 28°C, colonies were counted and assessed for color expression. Colonies without color expression were transferred to LB plates supplemented with nalidixic acid (to confirm *P. stewartii*) and gentamicin (to confirm retention of the plasmid).

### Plant inoculations with *Pseudomonas syringae*

Tomato seedlings (*Solanum lycopersicum* cv Moneymaker) were grown from seed and transplanted into 10 cm pots of Sunshine Mix #1 professional growing mix (SunGro Horticulture) in a temperature-controlled greenhouse maintained at 24-37°C with natural light. Plants were inoculated at 4 weeks old. Prior to inoculation, plants were brought into the laboratory and covered with plastic bags to increase humidity for 2 hours. Bacterial inoculum was prepared from *P. syringae* cultures grown on TSA plates for 24 hours. Bacterial cells were scraped off plates and suspended in 10 mM MgCl_2_ with 0.2% Tween 20 at a concentration of OD_600nm_ = 0.5. Plants were sprayed with bacterial inoculum to cover all leaf surfaces, and enclosed in plastic bags in the dark at 25°C for 24 hours. After the first 24 hours, plants were maintained indoors under fluorescent lighting at 25°C for five more days. At six days post-inoculation, one full leaf was collected from each plant for bacterial isolation. Leaves were weighed, then surface sterilized in 70% ethanol for 30 seconds, 20% bleach for 30 seconds, and then rinsed in sterile water twice for 2 minutes each. Then leaves were transferred to mesh grinding bags (Agdia) and crushed in 2 mL 1XPBS. Buffer containing crushed plant material was serially diluted and plated on TSA medium supplemented with rifampicin to select for *P. syringae*. Colonies were counted and assessed for color expression after 3 days incubation at 28°C.

## Results and Discussion

### Modification with chromoproteins produces plant pathogen strains that are visibly colored

Four species of plant pathogenic bacteria were modified with up to six different chromoproteins to produce a range of strains that are visibly colored for tracing and differentiation in plant disease research. In *X. fastidiosa*, chromoproteins were integrated into the chromosome because this is an efficient and stable method for gene insertion in this pathogen (44, 48, 49). Successful transformants expressing five different chromoproteins (red, orange, yellow, blue and purple) were obtained (Fig. 1) in *X. fastidiosa*. Several attempts to transform *X. fastidiosa* with the green chromoprotein failed, and since green is not very distinct from the yellow chromoprotein, it was omitted for *X. fastidiosa* experiments. The purple chromoprotein did not produce visible color expression in *X. fastidiosa* during normal growth on plates or in individual colonies, but can be seen in concentrated cell pellets (Supplemental Fig. S1). Although not as intense as when highly expressed in *E. coli*, the red, orange, yellow, and blue chromoproteins produced visible color in *X. fastidiosa* individual colonies at 10-14 days of growth, or in concentrated growth at 5-7 days (Fig. 1). Similar results were seen in both *X. fastidiosa* strains, Temecula-1 (Fig. 1C) and M23 (Fig. 1D) for all of the chromoproteins.

In *P. stewartii*, *P. syringae*, and *X. campestris*, chromoproteins were expressed on the broad host-range plasmid pBBR5pemIK (46). This plasmid vector is generally stable without antibiotic selection due to the PemI/PemK toxin-antitoxin system, and is highly versatile because it can be transformed into a wide range of bacterial species (46, 50). In *P. stewartii*, all six chromoprotein-modified strains were visibly colored after 3 days of growth and appeared to grow normally in culture (Fig. 1E). Due to naturally occurring yellow pigment in *P. stewartii* (51), yellow and green strains were more difficult to distinguish from wild type visually, but the color difference is still apparent if the strains are placed side by side and incubated for an additional 1-2 days. In *P. syringae*, color expression was visible in all five of the six chromoprotein-modified strains after 3 days of growth (Fig. 1F). The exception was the blue *P. syringae* strain which grew very slowly and had color expression that was unstable, resulting in reversion to wild type color. Colonies of the *P. syringae* blue strain which reverted back to wild type color retained resistance to gentamicin and were confirmed to still carry the pBBR5pemIK-BLUE plasmid. Sequencing of pBBR5pemIK-BLUE extracted from *P. syringae* colonies that lost color expression found an introduced internal stop codon in the chromoprotein gene Supplemental Figure S2. This suggests that the blue chromoprotein has negative effects in *P. syringae* which are overcome by mutation rather than loss of the plasmid.

The red, orange, yellow, green, and purple chromoproteins did not appear to have any negative impacts on growth of *P. syringae* and produced strong color expression. *Xanthomonas* species also produce pigments (52), and in *X. campestris* the naturally occurring yellow pigments obscured visible expression of some of the chromoproteins (Fig. 1G). In *X. campestris*, red, orange, and purple chromoprotein expression was visible after 3 days of growth, but yellow and green were difficult to distinguish from the wild type color, and blue did not appear to be expressed at all. Unlike in *P. syringae*, pBBR5pemIK-BLUE extracted from *X. campestris* was intact when sequenced. This suggests that the blue chromoprotein is not necessarily detrimental in *X. campestris*, just that the color is not visible.

Different results of chromoprotein expression across the four different plant pathogen species highlights the utility of having multiple different color options so that the best performing chromoproteins can be used depending on the study system. Overall, for each of the four species there were 4-6 different chromoproteins that produce highly visible colors growing in culture. These options could facilitate a wide range of experimental possibilities relating to strain-strain, or species-species interactions in vitro.

### Chromoprotein-modified *X. fastidiosa* strains have stable color expression during plant infection

Two strains of *X. fastidiosa* (Temecula-1 and M23) modified with four different chromoproteins (red, orange, yellow, and blue) were inoculated into grapevines to assess pathogenicity and stability of the color expression. Strains modified with the purple chromoprotein were not used for plant inoculations due to low color visibility observed in vitro. All eight *X. fastidiosa* strains (four in Temecula-1 background and four in M23 background) caused leaf scorch symptoms in susceptible grapevines starting at 8-10 weeks post-inoculation (Fig. 2, Fig. S3). Although there was no obvious difference in ability of the chromoprotein-modified strains to cause disease symptoms, larger samples sizes and more extensive inoculation experiments would be necessary to fully evaluate whether there are any quantitative differences in virulence of these strains compared with the wild type parent strains.

**Figure 2.**
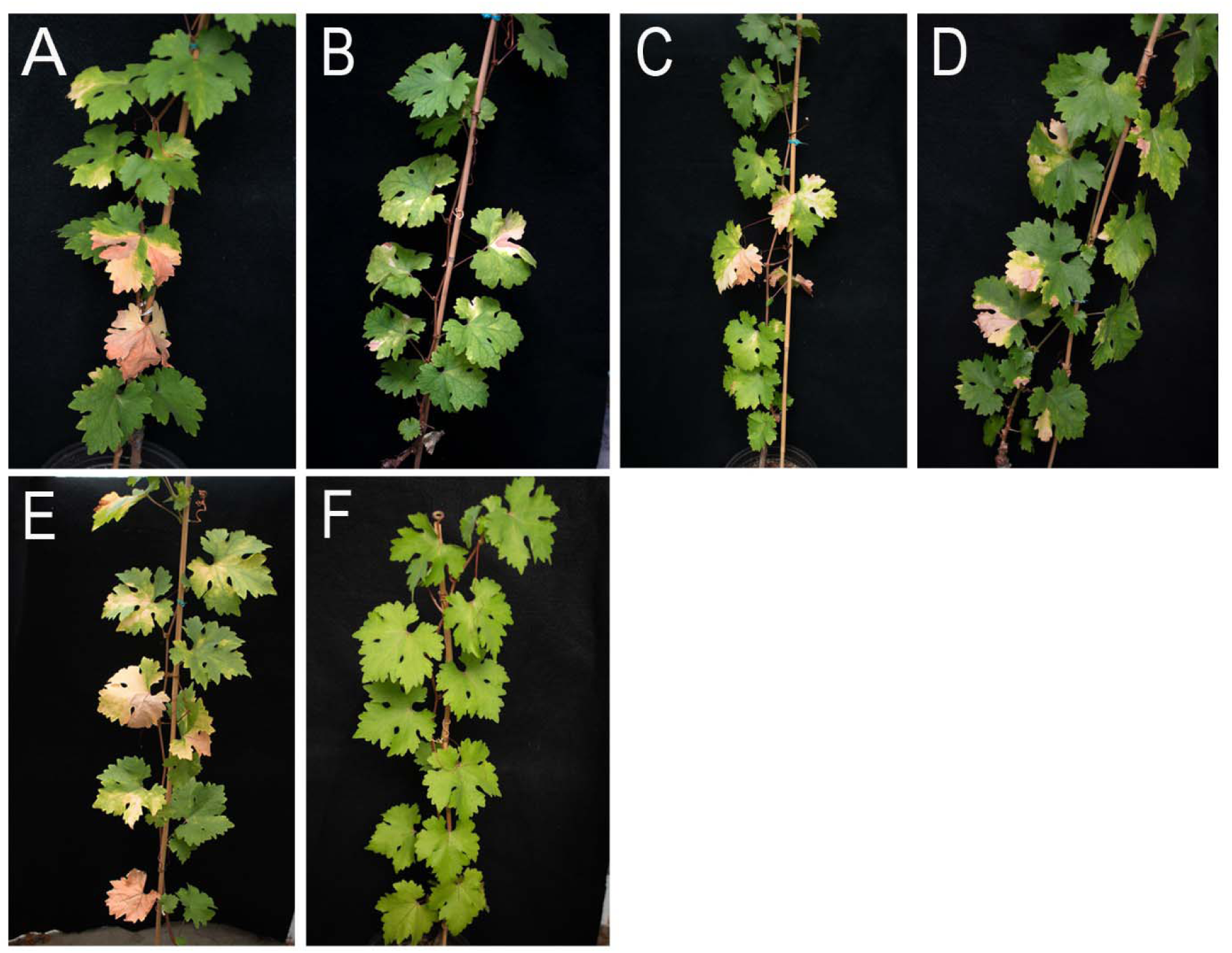
Leaf scorch symptoms in grapevines inoculated with chromoprotein modified *Xylella fastidiosa* strains in the M23 background. Plants were inoculated using the pin-prick inoculation protocol and maintained in a greenhouse for 12 weeks. Leaf scorch disease symptoms were visible on all inoculated plants beginning at 8 weeks post-inoculation for plants inoculated with the following chromoprotein-modified strains: A) red, B) orange, C) yellow, D) blue, E) wild type M23, F) mock-inoculated negative control.

Bacterial quantity isolated from petioles of symptomatic leaves were similar across plants inoculated with chromoprotein-modified strains compared with the wild type strains (Fig. 3), and none of the modified strains had significantly different bacterial quantities than the wild type parent strain based on ANOVA and post-hoc means comparison (Tukey, p<0.05, Fig. 3A). Color expression was retained in all isolated colonies from all samples (Fig. 3B), indicating that the chromoprotein modifications are stable during plant infection. These results show that chromoprotein-modified *X. fastidiosa* strains can be used for in planta experiments for pathogen tracking.

**Figure 3.**
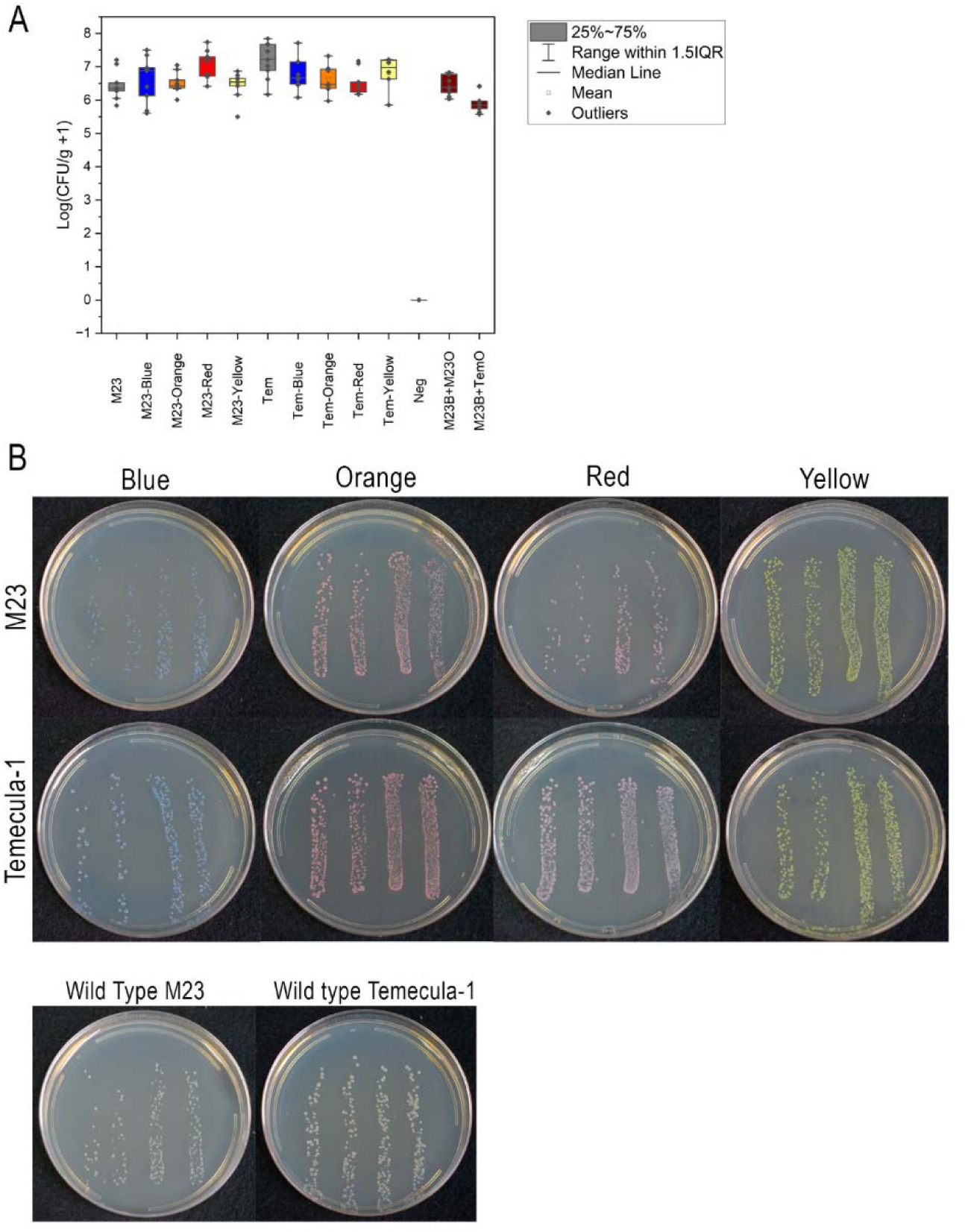
Bacterial populations isolated from grapevines inoculated with chromoprotein-modified *Xylella fastidiosa* strains. Bacteria were isolated from each inoculated plant once the plant reached a disease score of 2-3 on a 0-5 scale (8-10 weeks post-inoculation). Three petioles were collected from symptomatic leaves on each plant, surface sterilized, and ground in 1XPBS. Serial dilutions of the ground plant material were plated on PD3 plates and incubated at 28°C for 14 days for colonies to be large enough to count and for color expression to be visible. A) Colony forming units (CFU) were normalized to plant tissue weight and log transformed for plotting. Bacterial quantities from plants inoculated with chromoprotein-modified strains were not significantly different from plants inoculated with the wild type strains based on ANOVA and post-hoc means comparison testing (Tukey, p<0.05). B) Plates were photographed after 14 days of incubation to visualize colors.

### Chromoprotein-modified *Xylella fastidiosa* strains can be tracked in co-inoculation experiments

Plants inoculated with a 1:1 mix of M23-Blue and M23-Orange or with a 1:1 mix of M23-Blue and Temecula-Orange were evaluated the same as described above for the plants inoculated with single strains. Total bacterial quantities isolated from plants inoculated with two different strains were not significantly different from bacterial quantities in wild type inoculated plants (Fig. 3A). Proportion of each strain varied between individual samples (Fig. 4A), but there was no significant difference in quantity of blue and orange chromoprotein-modified strains overall, regardless of the parent strain background (Fig. 4B). The chromoprotein-modified strains can be easily differentiated visually upon isolation from plant tissues (Fig. 4C). This suggests that in addition to being stable during plant infection, the chromoprotein-modified *X. fastidiosa* strains could be used to compare spread of different strains within a plant, or to evaluate competition of knockout mutants. The two wild type backgrounds used in these experiments, Temecula-1 and M23, are both *X. fastidiosa* subsp. *fastidiosa* and belong to the same sequence type (ST1). As both these strains are virulent in *V. vinifera* grapevines it is not expected that there would be a significant competitive advantage between them. However, chromoprotein modification could be used to compare *X. fastidiosa* strains of more diverse genetic backgrounds, or in a wider range of host plants to better understand competition and selection dynamics in this pathogen.

**Figure 4.**
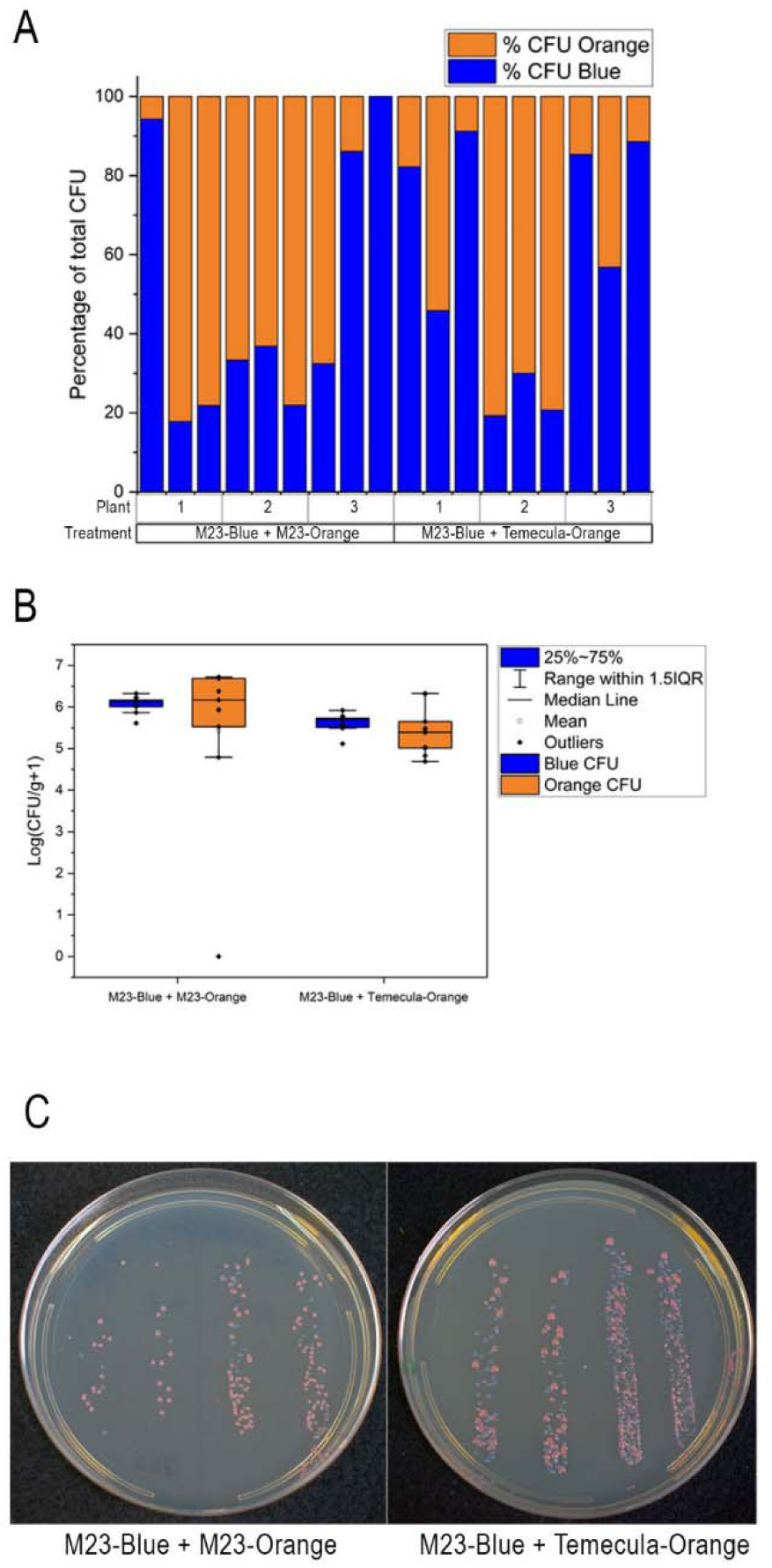
Co-inoculation with chromoprotein-modified *Xylella fastidiosa* strains of two different colors. Grapevines were inoculated with a 1:1 mix of M23-Blue and M23-Orange, or M23-Blue and Temecula-Orange strains. After 10 weeks post-inoculation, three petiole samples were collected from symptomatic leaves on each plant for bacterial isolation. Proportion of the different chromoprotein-modified strains was compared for each sample (A), as well as relative quantities of the different strains in each treatment overall (B). Quantity of blue vs orange colonies in each treatment was compared using a Student’s t-test (p<0.05) and was not significantly different regardless of the strain background. Comparisons of blue vs orange chromoprotein-modified strains was done by visual evaluation of the colonies on plates (C).

### Stability of plasmid-based chromoprotein expression in *Pantoea stewartii* during plant infection varies based on chromoprotein and experimental conditions

Inoculation of chromoprotein-modified *P. stewartii* strains in susceptible sweet corn seedlings produced water-soaked lesions and significant bacterial colonization (Fig. 5). Although lesion severity was reduced in all modified strains including the empty vector control (Fig. 5A), bacterial quantities isolated from infected leaf sections were not significantly different from while type population levels (Fig. 5B). In these experiments, *P. stewartii* was inoculated as a cell suspension placed in the seedling whorl which typically leads to water-soaked lesion formation and entry of the bacterium into the apoplast (53). Other inoculation methods intended to more closely mimic entry of *P. stewartii* into the xylem vessels via insect feeding use wounding of the stems for direct uptake of the bacterium into the xylem (54). It is possible that disease development from the chromoprotein-modified strains would vary depending on the inoculation method, as this is the case for some deletion mutants (55). It is also possible that disease severity caused by the chromoprotein-modified strains would be impacted by environmental conditions, plant inoculations were conducted in a greenhouse without humidity control in this case. Regardless, the chromoprotein-modified *P. stewartii* retained some ability to produce water-soaked lesions and replicate to high levels in plant tissue. However, chromoprotein-expression in colonies isolated from the plant varied by strain and between experiments suggesting these constructs are not completely stable during plant infection (Fig. 6). Inoculations experiments were performed twice, and in the first experiment the percentage of color-expressing colonies was over 80-90% for most of the strains except for the blue strain which had a significantly lower percentage (Fig. 6A). In the second experiment, the percentage of color-expressing colonies was lower for all strains at between 40-60% for all except for the blue strain which had very few colored colonies at all (Fig. 6B). Blue colonies also had a mucoid phenotype indicative of exopolysaccharide (EPS) overexpression which was not observed in any of the other strains (Fig. 6C). EPS is an important virulence factor for *P. stewartii*, and the timing and amount of EPS produced is tightly regulated during the infection process (56, 57). Disruption to normal EPS production could be why the blue chromoprotein was significantly less stable than the others. It is not clear why expression of the blue chromoprotein in *P. stewartii* would change EPS production, but it is a useful consideration for future use of these constructs for plant disease assays. For all the strains, colonies that did not have visible color expression were screened on selective media containing nalidixic acid (to confirm *P. stewartii* and rule out contamination), and gentamicin to evaluate plasmid retention. At least 50 non-colored colonies were screened for each strain, and for the chromoprotein-modified strains less than 5% of these colonies were resistant to gentamicin suggesting that loss of color expression is primarily due to loss of the plasmid. Over 200 colonies were screened from the empty vector control, and of these 100% retained gentamicin resistance which implies that the pBBR5pemIK plasmid is not inherently unstable in *P. stewartii* during plant infection, but rather that chromoprotein expression may exacerbate any fitness effects in planta.

**Figure 5.**
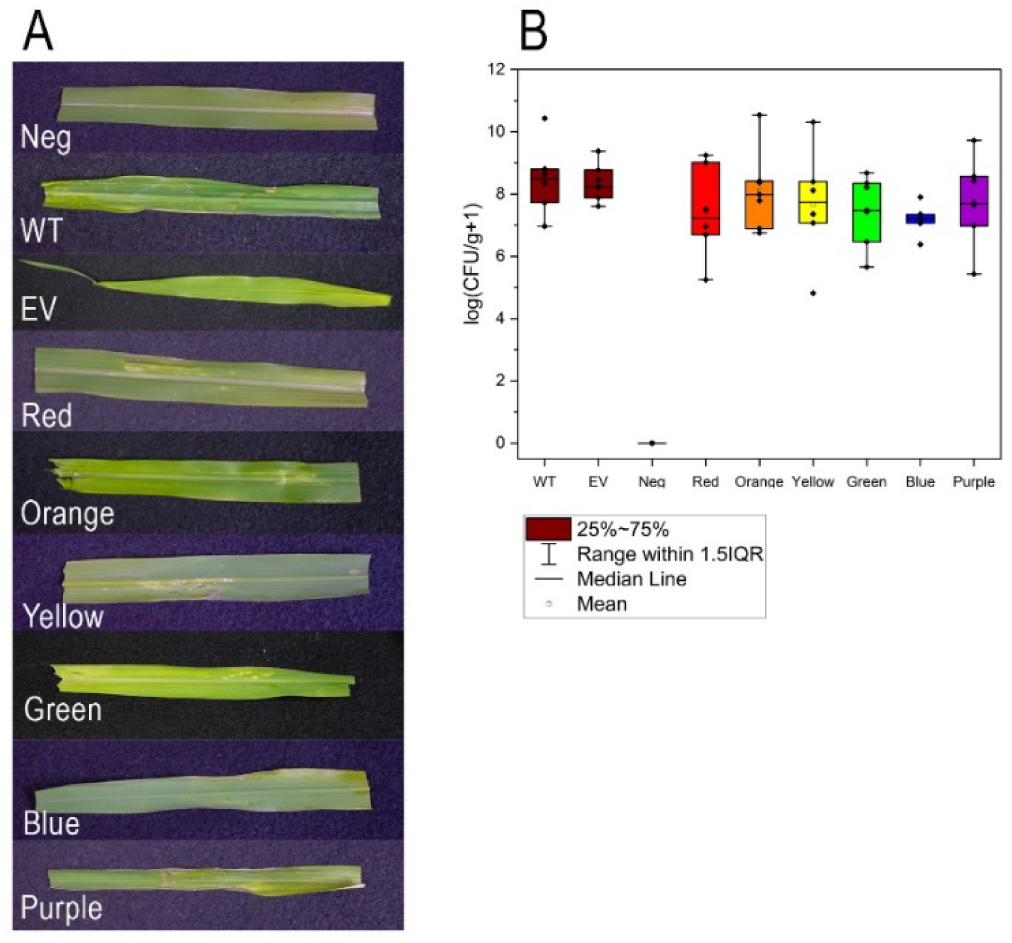
Chromoprotein-modified *Pantoea stewartii* infection in sweet corn seedlings. Plants were inoculated with bacterial suspension into the seedling leaf whorl and evaluated for disease symptoms (A) and bacterial colonization (B) after 4 days. All inoculated plants had some degree of water-soaked lesions, but all of the modified strains including the empty vector control had less extensive lesions than the wild type. Total bacterial quantities isolated from a 10 cm leaf section were not significantly different in any of the inoculation treatments based on ANOVA and post-hoc means comparison (Tukey, p<0.05). Bacterial quantities are represented as log-transformed CFU normalized to plant tissue weight. WT = wild type, EV = empty vector control, Neg = mock-inoculated negative control.

**Figure 6.**
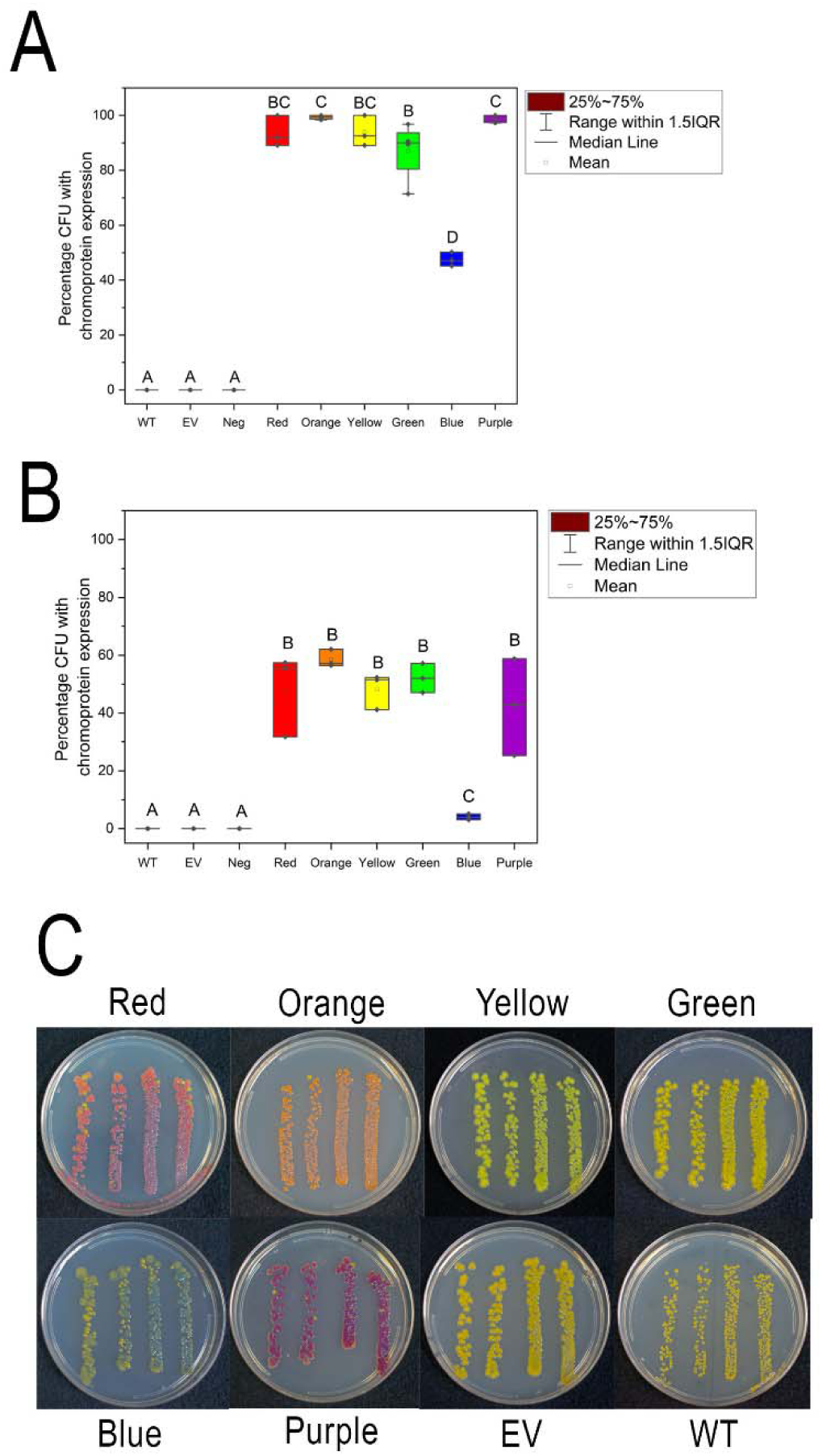
Stability of plasmid-based expression of chromoproteins in *Pantoea stewartii* during plant infection. Populations of *P. stewartii* isolated from plant tissue at 4 days post-inoculation did not uniformly retain color expression. Inoculation experiments were performed twice, and in the first experiment (A) percentage of color-expressing colonies was generally higher than in the second experiment (B). There were also significant differences between the different chromoprotein-modified strains as evaluated by ANOVA and post-hoc means comparison testing (Tukey, p<0.05). Bars marked with the same letter are not significantly different from each other. Color expression was most visible for counting after 4 days of growth on plates (C). Photographs are from experiment (A). WT = wild type, EV = empty vector control, Neg = mock-inoculated negative control.

Again, the environmental conditions during infection could be a factor in stability of P. stewartii chromoprotein expression during plant infection. Since these experiments were performed in a greenhouse, the temperatures could vary within the range of 28-37°C which might impact plasmid loss, and stability of the constructs might be better under more optimal disease conditions. It is also possible that chromoprotein-modified *P. stewartii* strains created by chromosomal insertion rather than plasmid expression would be a significant improvement. Despite the instability in planta, these experiments provide some useful insight into which chromoproteins have potential for experimental use in the *Pantoea* genus, and lay the groundwork for optimization for specific research scenarios.

Stability of plasmid-based chromoprotein expression in *Pseudomonas syringae* during plant infection varies based on chromoprotein. Tomato plants inoculated with chromoprotein-modified *P. syringae* strains all developed foliar disease symptoms similar to the wild type strain (Fig. 7A). All of the strains tested also were able to replicate in the plant to population levels that were not significantly different from the wild type in six days after inoculation (Fig. 7B). Due to the instability of the blue chromoprotein construct in *P. syringae* in vitro, the blue strain was not tested in plants. Retention of chromoprotein expression in *P. syringae* isolated from inoculated plants varied depending on the specific color construct (Fig. 8), with the highest retention in the yellow strain (80-90%) and the lowest in the red strain (30-50%). Orange and purple strains had intermediate retention of color expression in the isolated colonies. For the green strain, color expression was not intense enough to visually differentiate colonies with and without color expression, so this strain could not be evaluated for stability this way. An additional challenge in *P. syringae* is that color expression in single colonies is not highly visible until 3-4 days of growth on plates, at which point colonies are very large (Fig. 8B). As in *P. stewartii*, colonies without color expression were screened for gentamicin resistance to evaluate plasmid loss. Between 50 and 100 non-colored colonies were screened per strain, and 100% of these were no longer resistant to gentamicin indicated plasmid loss. Of the over 200 colonies screened for the empty vector strain, all were still gentamicin resistant suggesting that the pBBR5pemIK plasmid is not inherently unstable in *P. syringae* during plant infection, but rather that chromoprotein expression drives the plasmid loss. In *P. syringae* infection via spray inoculation, it is possible that insertion of chromoprotein genes into the chromosome could be more effective and stable than plasmid-based expression. Due to the variability between the different strains however, these results provide some insight into which chromoproteins might be better suited to expression in *P. syringae* and which ones might be less effective for plant disease assays in this species.

**Figure 7.**
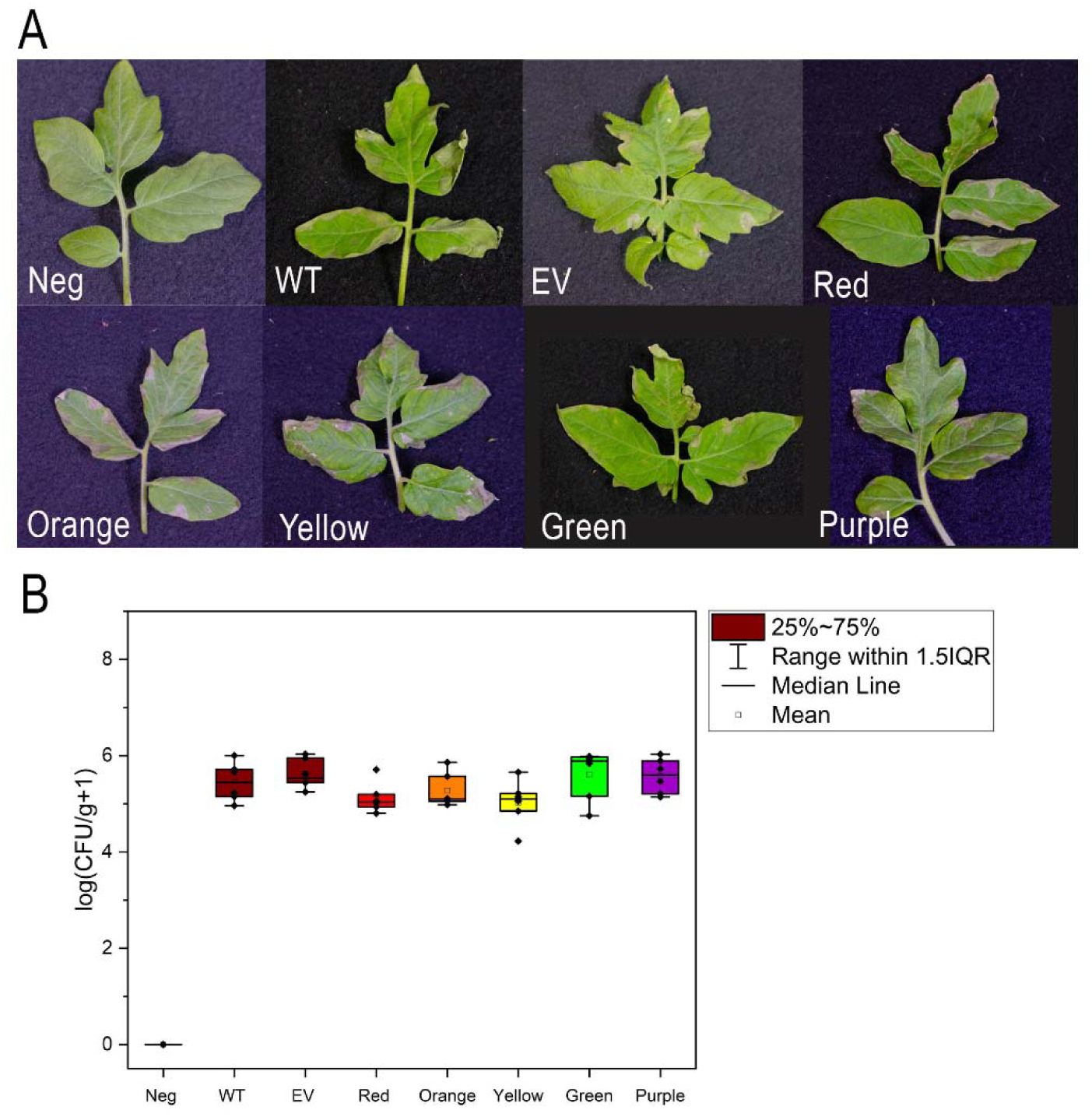
Chromoprotein-modified *Pseudomonas syringae* infection in tomato plants. Plants were inoculated by spraying with bacterial suspension and evaluated for disease symptoms (A) and bacterial colonization (B) after 6 days. All inoculated plants had some disease lesions on the leaves that were similar in severity across treatments. Total bacterial quantities isolated from a single leaf were not significantly different in any of the inoculation treatments based on ANOVA and post-hoc means comparison (Tukey, p<0.05). Bacterial quantities are represented as log-transformed CFU normalized to plant tissue weight. WT = wild type, EV = empty vector control, Neg = mock-inoculated negative control.

**Figure 8.**
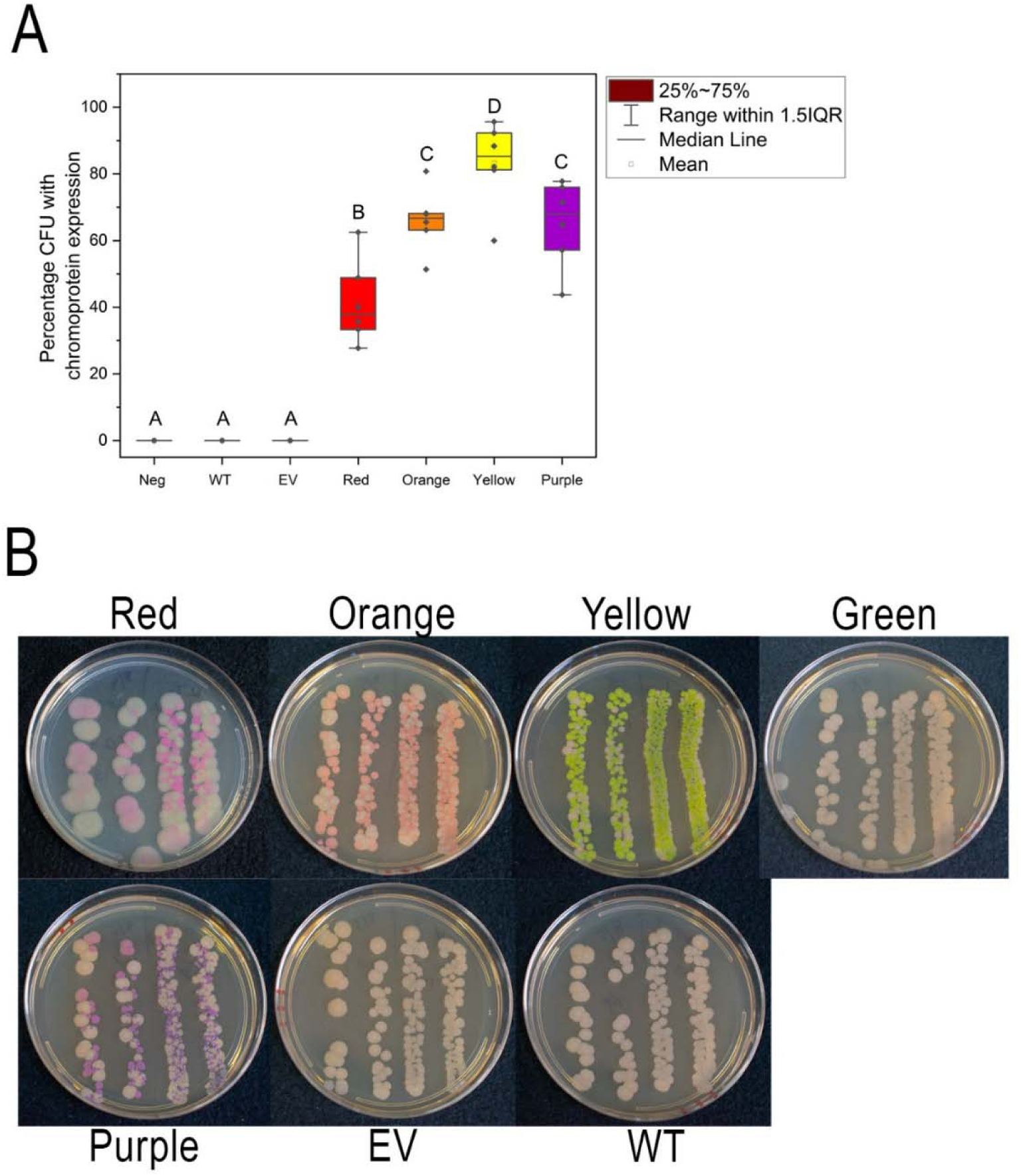
Stability of plasmid-based expression of chromoproteins in *Pseudomonas syringae* during plant infection. Populations of *P. syringae* isolated from plant tissue at 6 days post-inoculation did not uniformly retain color expression. Percentage of color-expressing colonies varied between strains (A). Bars marked with different letters are significantly different based on ANOVA and post-hoc means comparison testing (Tukey, p<0.05). The green chromoprotein strain was not possible to evaluate for percentage of colonies with color expression due to the color being too similar to wild type after extended growth. Color expression was most visible after 4 days of growth on plates (B). Experiment was performed twice and there was no significant difference between percentage of color-expressing colonies between the two experimental replicates. WT = wild type, EV = empty vector control.

## Conclusions

This study evaluated a set of six different chromoproteins in four different plant pathogenic bacterial species for creating marked strains for tracking in disease assays and transmission experiments. Several of these markers were highly successful for creating intensely colored bacterial strains without a significant burden on growth or pathogenicity. Interestingly, the results varied between bacterial species and chromoprotein used, highlighting the need for testing these more broadly and optimizing markers for a specific microbial-host system. The chromosomally inserted chromoprotein constructs in *X. fastidiosa* were very stable during plant infection, in contrast with plasmid-based expression that was tested in *P. stewartii* and *P. syringae*. However, even within the plasmid-based marked strains there were differences in stability and growth impacts depending on the bacterial species and specific chromoprotein that was used. There were also striking differences in color intensity between species. Purple for example, was one of the brightest colors in *P. stewartii*, *P. syringae*, and *X. campestris*, but was hardly visible in *X. fastidiosa*. Blue was unstable in *P. syringae* and *X. campestris*, but highly visible and stable in *X. fastidiosa*. The impact of naturally occurring pigments in the different bacterial species on color visibility also varied. Although *P. stewartii* naturally produces a yellow pigment, most of the colors were still highly visible. In *X. campestris* on the other hand, most of the colors were difficult to see in colonies due to the pigmentation and EPS production. Difficulty distinguishing most of the colors is why *X. campestris* strains were not tested in plant assays. Overall, these results provide some useful information for improvement and optimization of marker strains for plant disease research, building on prior work that identified and adapted these chromoproteins for bacterial expression (17, 24, 58). Additionally, the successful cases demonstrated here, such as in *X. fastidiosa*, will serve as a resource for the research community to facilitate investigation of pathogen dynamics in complex host and environmental scenarios.

## Data availability

Sequences of plasmids created in this study are available in NCBI as accession numbers: PV932843-PV932847 and PV983734-PV983739. All plasmids, constructs, and bacterial strains produced in this study are available from the corresponding author on request.

## Acknowledgements

Funding support was from the United States Department of Agriculture, Agricultural Research Service appropriated project #2034-22000-014-000-D. Mention of trade names or commercial products in this publication is solely for the purpose of providing specific information and does not imply recommendation or endorsement by the U.S. Department of Agriculture.

