## Supplemental Figure S1-S4 for "Chromoprotein-modified plant pathogenic bacteria: tools for experimental tracking and visualization"

**Supplemental Figures and Tables**

Supplemental Figure S1. Concentrated cell pellet of chromoprotein-modified *Xylella fastidiosa* strains.

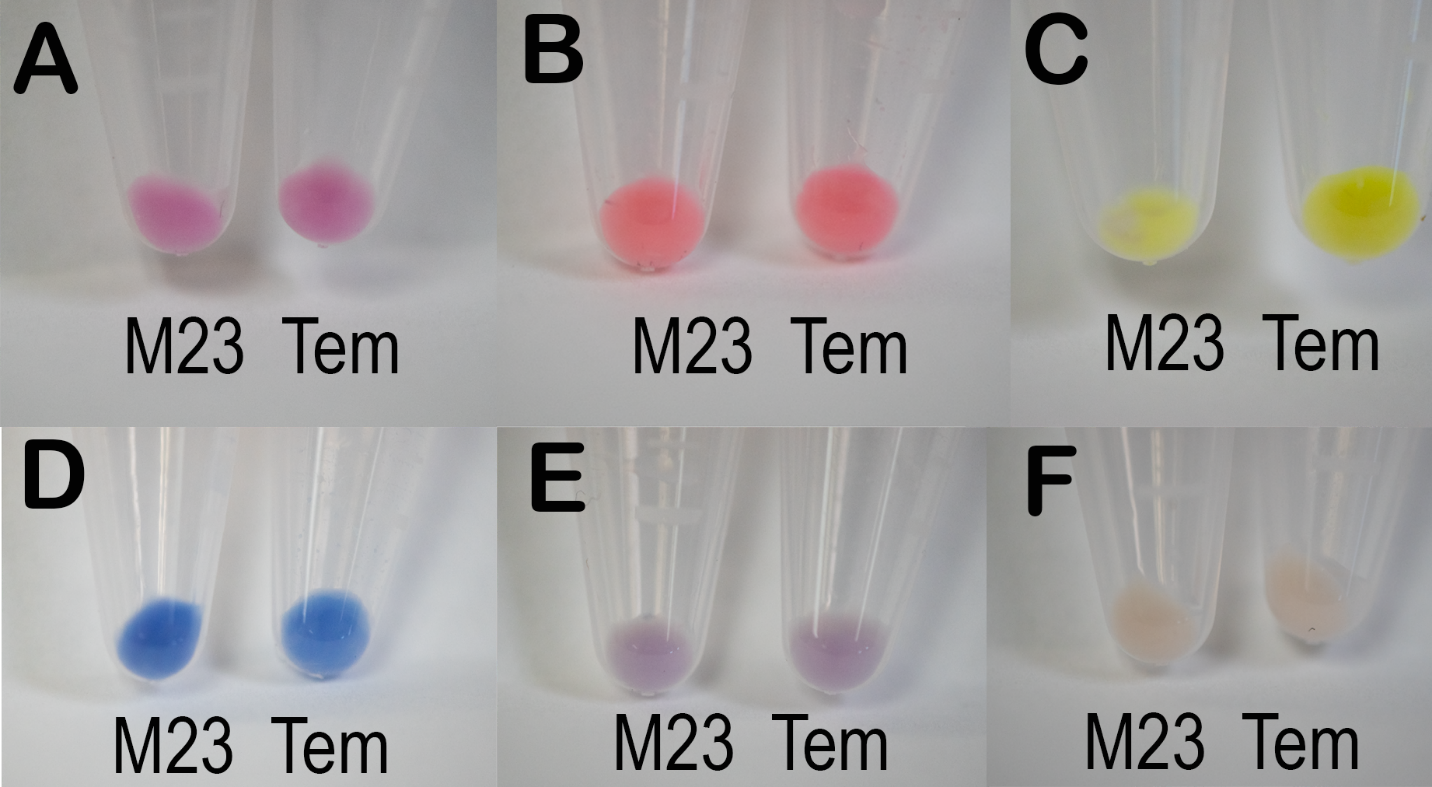

Cells of *Xylella fastidiosa* strains M23 and Temecula-1 (Tem) modified with A) red, B) orange, C) yellow, D) blue, and E) purple chromoproteins compared with F) wild type cell color.

**Supplemental Figure S2.** Mutated sequence of pBBR5pemIK-BLUE extracted from *Pseudomonas syringae* after loss of color expression.

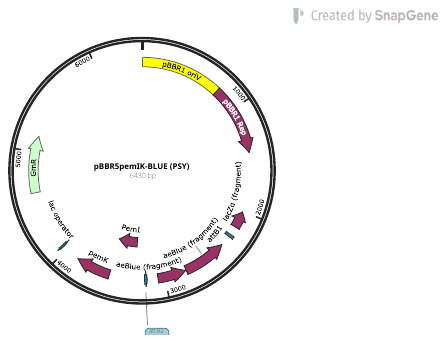

**Supplemental Figure S3. Leaf scorch symptoms in grapevines inoculated with chromoprotein-modified Xylella fastidiosa strains in the Temecula-1 background.** A) red, B) orange, C) yellow, D) blue, E) wild type Temecula-1, F) mock-inoculated negative control.

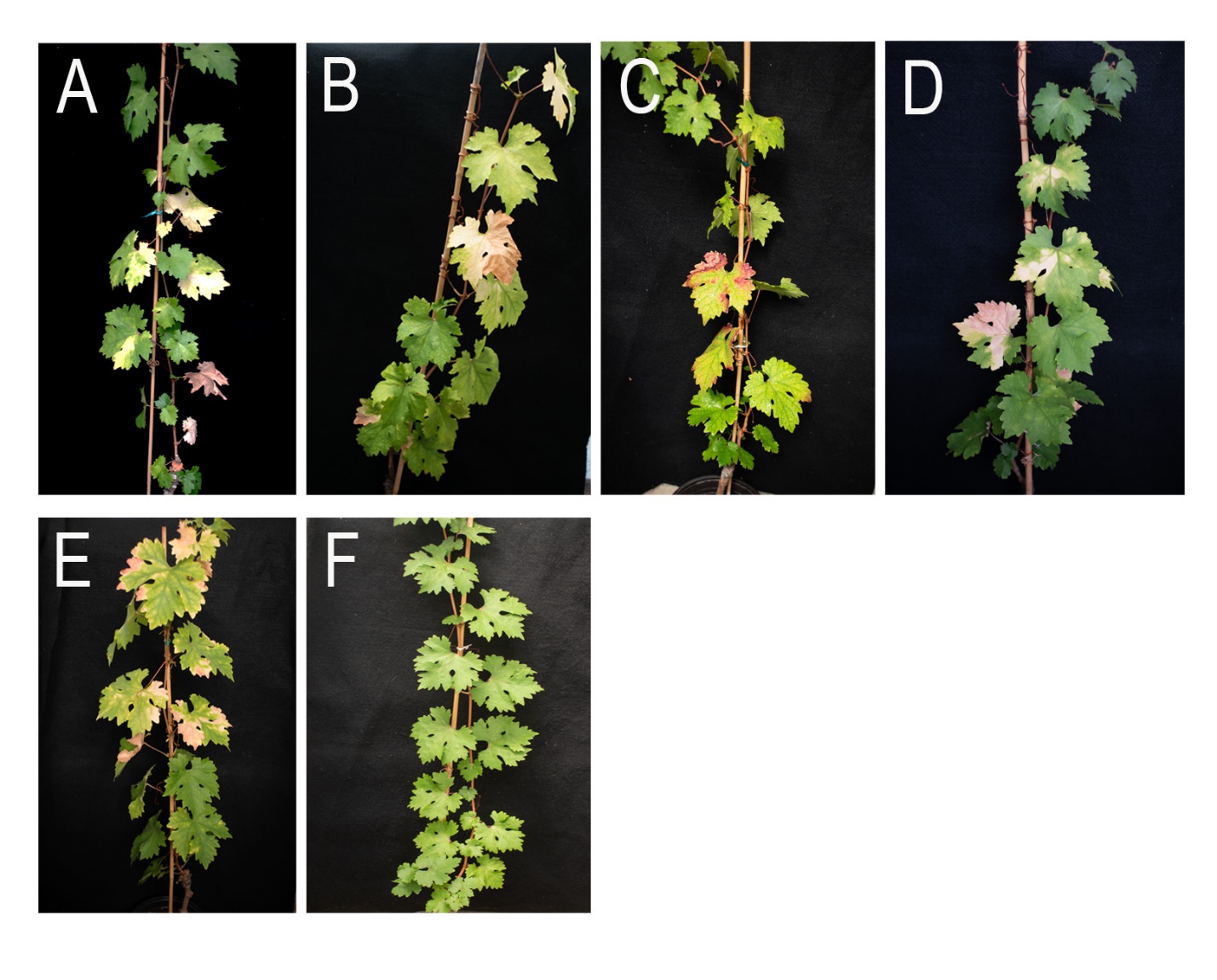
